# Compressing Streams of Phylogenetic Trees

**DOI:** 10.1101/440644

**Authors:** Axel Trefzer, Alexandros Stamatakis

**Affiliations:** Karlsruhe Institute of Technology, Karslruhe, Germany; Heidelberg Institute for Theoretical Studies, Heidelberg, Germany

## Abstract

Bayesian Markov-Chain Monte Carlo (MCMC) methods for phylogenetic tree inference, that is, inference of the evolutionary history of distinct species using their molecular sequence data, typically generate large sets of phylogenetic trees. The trees generated by the MCMC procedure are samples of the posterior probability distribution that MCMC methods approximate. Thus, they generate a stream of correlated binary trees that need to be stored. Here, we adapt state-of-the art algorithms for binary tree compression to phylogenetic tree data streams and extend them to also store the required meta-data. On a phylogenetic tree stream containing 1, 000 trees with 500 leaves including branch length values, we achieve a compression rate of 5.4 compared to the uncompressed tree files and of 1.8 compared to bzip2-compressed tree files. For compressing the same trees, but without branch length values, our compression method is approximately an order of magnitude better than bzip2. A prototype implementation is available at https://github.com/axeltref/tree-compression.git.

## Introduction

Phylogenetic trees, or phylogenies, for short, are unrooted, strictly binary, leaf labeled trees. They represent the evolutionary history of a set of *n* molecular sequences (e.g., DNA sequences) that correspond to the *n* distinct species under study. The leaves represent extant species, while the inner nodes represent hypothetical common ancestors. Henceforth, we refer to species/leaves as taxa. Phylogenies have various important applications in biological and medical research (e.g., [1]).

At present, likelihood-based statistical models of evolution are widely used to reconstruct such trees, either under the Maximum Likelihood (ML) criterion or using MCMC-based Bayesian Inference (BI) methods (e.g., [2]). Due to the super-exponential increase in the number of possible trees as a function of the number of taxa, ML-based phylogenetic inference is known to be NP-hard [3].

BI methods strive to compute the posterior probability distribution of phylogenetic trees. Here, the computational complexity is determined by the marginal probability term, whose exact calculation requires evaluating the likelihood of *all* possible tree topologies. Thus, MCMC methods are deployed to approximate the posterior. In general, the MCMC procedure is run for several million generations, and tree samples are typically written to file every 1, 000 generations. Therefore, MCMC runs,especially on datasets with a large number of taxa *n*, can generate tree files of substantial size. To alleviate this problem we investigate if state-of-the art algorithms for compressing binary trees can be applied and adapted to phylogenetic trees.

In the following we outline the main challenges. First, the trees contain branch lengths that represent the genetic distance between two nodes in the tree. Second, the tree samples can be regarded as a data stream. Third, successive tree samples are generally highly correlated. Thus, the tree topology and branch lengths between sample *i* and sample *i*+1 do typically not differ substantially. We incorporate all these characteristics in our compression algorithm to improve compression of phylogenetic tree streams.

The remainder of this paper is organized as follows: We first review related work on general tree compression and phylogenetic tree compression. Then, we describe our compression algorithm and discuss some implementation details. Thereafter, we present and discuss our results on empirical datasets. Finally, we provide a conclusion and discuss directions of future work.

## Related Work

### General binary tree compression

The topology of a *n*-node tree can be encoded in only 2*n* bits by traversing it in depth-first order and appending an opening parenthesis when entering a node and a closing parenthesis when leaving it. This encoding is called balanced parentheses (BP) as a balanced number of opening and closing parentheses encodes each subtree. While this encoding requires substantially less space than a pointer-based tree representation (requiring *O*(*n* log *n*) bits), navigating the tree (e.g., finding a parent or the *i*-th child) requires a costly sequential decompression of parts of the tree. Jacobson [4] introduced constant time basic navigation operations via an additional sub-linear index of size *o*(*n*) bits. This result was the starting point of succinct data structures; they encode instances of objects in space that is close to the information theoretic minimum for distinguishing among all possible instances. For trees, this bound is 2*n –*Θ(*logn*) bits. Besides BP, there are two other popular succinct representations [4, 5] which offer different trade-offs for tree operations.

### Compressing phylogenies

We are not aware of previous work on compressing streams of correlated phylogenies. Ane and Sanderson [6] presented work on compressing phylogenies in conjunction with the associated sequence data. Matthews *et al.* [7] presented a method for compressing phylogenetic tree topologies, but without taking into account branch lengths. However, branch lengths constitute vital information for the majority of post-analysis tasks (e.g., for estimating divergence times or delimiting species).

## Compression Algorithm

Before describing our two compression algorithms, we outline the calculation of the Robinson-Foulds metric (RF metric[8]) for comparing phylogenetic trees with the same taxon set. The RF metric is a prerequisite for our more involved compression scheme.

### The Robinson Foulds Metric

The RF metric relies on the property that each edge/branch of a binary phylogeny splits the taxon set into two partitions: removing the edge generates two subtrees, each corresponding to one partition of the taxa. We call such splits bipartitions. Bipartitions at edges leading to leaves are trivial as they are present in any tree containing the taxon set. Thus, we focus on non-trivial bipartitions, that is, splits at inner edges of the tree. The set of *all* non-trivial biparitions of a tree is mathematically equivalent to the tree representation. The sets of non-trivial bipartitions of two trees can be used to compare them. The RF distance between two trees is defined as the number of bipartitions that are unique to one of the two trees.

The RF metric does not only provide a distance measure between two trees, but also allows for transforming one tree into another using appropriate tree edit operations. A so-called strict *consensus tree* can be computed, that only contains edges that are present in the bipartition sets of *both* trees. Such a strict consensus tree does not need (and is typically not) binary any more. As the strict consensus tree contains all edges that are common to both trees, our key idea to compress successive tree samples *i* and *i* + 1 from a phylogenetic tree stream is as follows: Contract edges in the tree *i* to transform it into the consensus tree, and then extend the consensus tree again to obtain tree *i* + 1. To perform this transformation, only the contraction operations for tree *i* and the expansion operations of the consensus tree (for constructing tree *i* + 1) have to be stored.

We show an example RF distance calculation for two trees with identical taxon sets in Figure 1. Non-RF branches, that is, branches that occur in both trees are shown in red. RF branches, that is, branches that are unique to one of the two trees are shown in green. We also display the resulting consensus tree.

**Figure 1:**
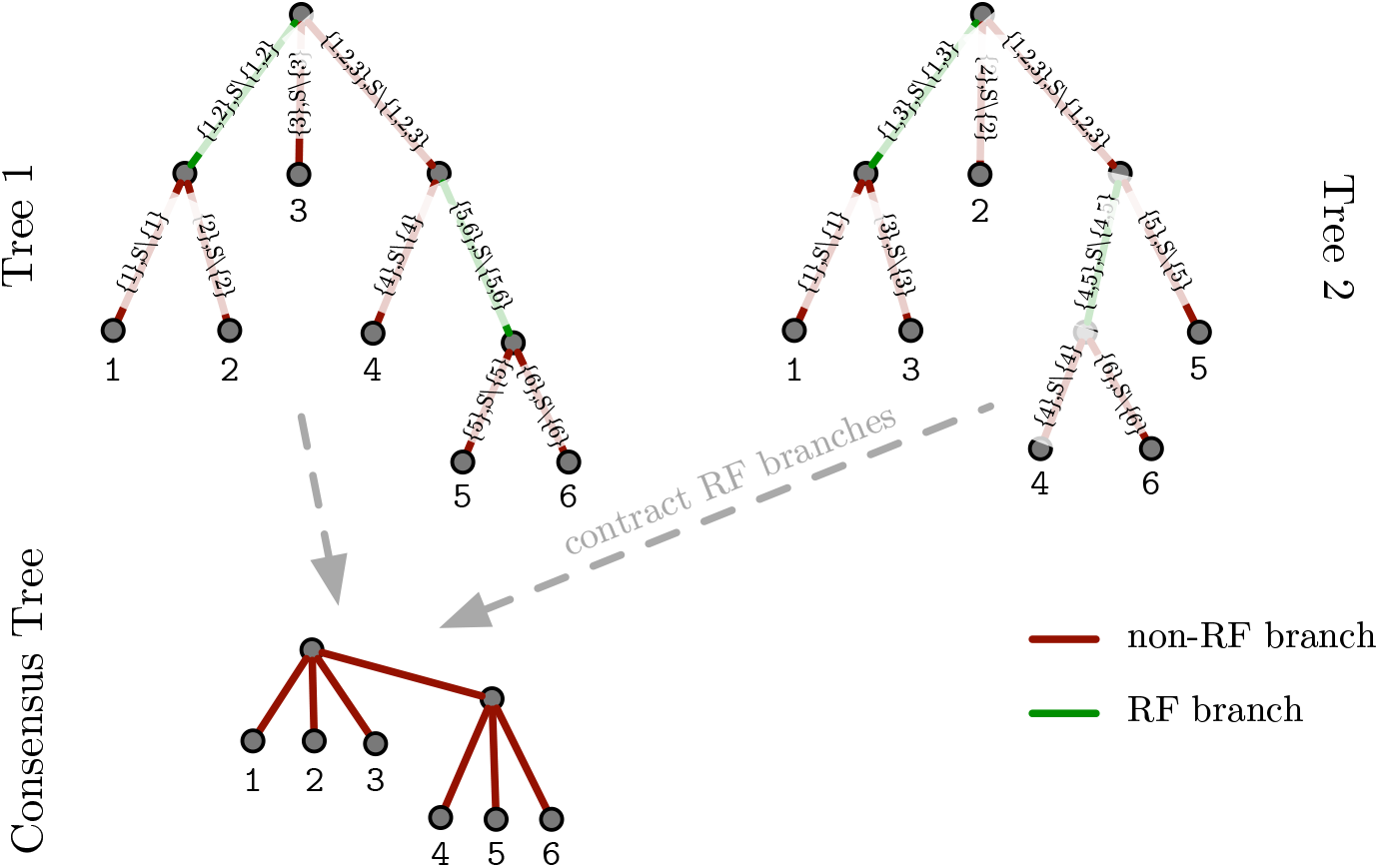
RF distance between two trees and consensus tree below.

#### Compression Algorithms

We implement two distinct compression methods: a simple compression (SC) and a RF-based compression (RFC). Both methods can be applied to phylogenies whose taxon names have been mapped to unique integer identifiers. The SC method can be applied to a single tree and compresses its topology and branch lengths. The RFC method compresses a tree pair by storing and compressing edit operations between the two trees. As the RFC can only be applied to tree pairs, the SC is useful for compressing the first tree sample of a phylogenetic tree stream. All successive trees of that stream can then be handled via RFC. Moreover, the SC of a tree is sometimes more efficient than the RFC. This is the case when the RF distance between two consecutive trees is large (i.e., substantial topological changes occurred). This is likely to occur during the burn-in phase of MCMC analyses where most topological alteration proposals are likely to be accepted as phylogenetic BI implementations typical start sampling from a random tree. The SC therefore yields a guaranteed upper bound for the space required to store a single tree. We describe the SC and RFC in more detail

### Simple Compression

The SC compression scheme can be applied to *any* phylogeny. It is thus not limited to compressing tree samples of MCMC runs. The SC deploys three vector structures for compression:

- a bit vector storing the topology of the tree in BP encoding [4]
- an integer vector storing the permutation of the taxa at the leaves
- a double vector storing the branch lengths

Since all topologies in a MCMC tree stream comprise the same taxa, they also have an identical number of branches. Therefore, the vectors storing the topology and the leave permutation have constant size. However, the size of the branch length vector *can* vary, as we do compress it (see paragraph on branch length compression for details). Overall, the variance of the SC space requirements is low over a phylogenetic tree stream (see experimental results).

To calculate the SC, we first place a root into the tree we intend to compress. Without loss of generality, we root the unrooted phylogeny on the external branch leading to taxon identifier 1. This rooting method ensures a consistent representation of all trees in a stream. Then, we reorder the tree such that, when traversing it via a depth-first search (DFS), we always visit the subtree containing the smallest taxon number first. On this reordered tree (note that phylogenies are unordered trees), we now calculate the BP representation and the leaf permutation induced by a DFS traversal starting at the root. We also store the branch lengths in DFS order. An example showing the tree reordering and the respective data structures is provided in Figure 2.

**Figure 2:**
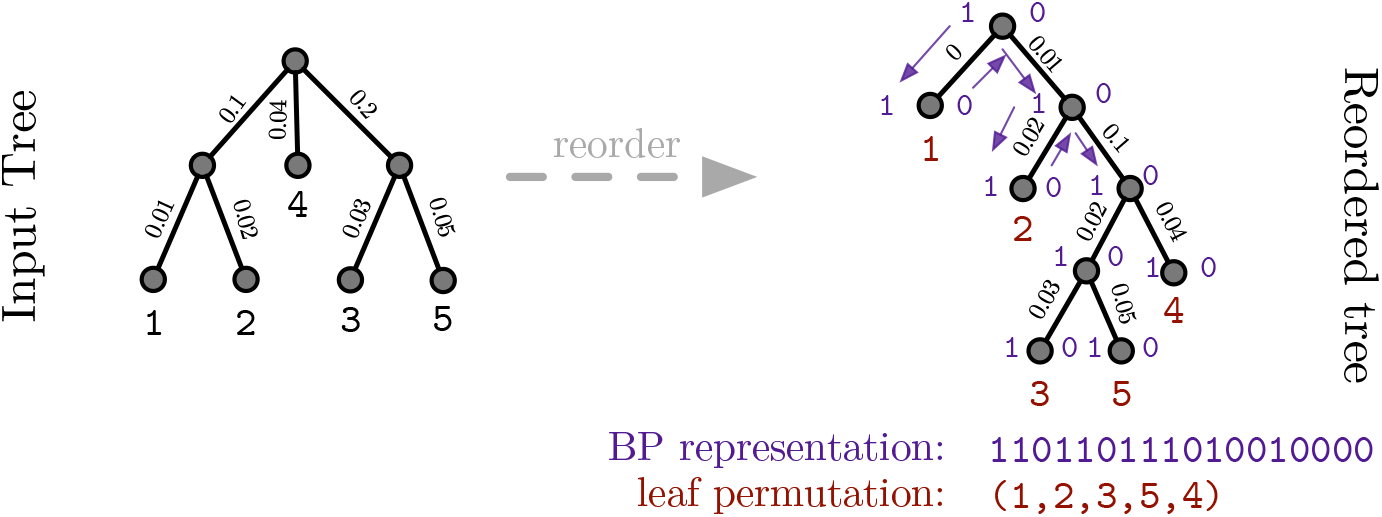
SC operation example

The decompression is straight-forward: we build the tree using the BP encoding, assign the respective taxon identifiers, and branch lengths during tree construction.

### Robinson-Foulds Compression

As mentioned before, the RFC operates on tree pairs, typically two consecutive tree samples from a MCMC implementation. The compression relies on the RF distance between the two trees. As for SC, we first transform both trees into their rooted and ordered representation. Then, we calculate the RF distance between them. Based on the RF distance, we classify the edges as:

1. branches/edges, that contribute to the RF distance (RF branches), that is, they do not form part of a strict consensus tree
2. branches/edges, that do not contribute to the RF distance (non-RF branches), that is, branches, that are shared among both trees and will hence be included in a strict consensus tree

Our main idea is to only store the edit operations that we need to apply to the first tree to obtain the second tree. These edit operations comprise the contraction of RF branches in the first tree (leading to the strict consensus tree) and the expansion of the strict consensus tree by expanding the RF branches from the second tree. An example of these operations and the associated data structures is provided in Figure 3.

**Figure 3:**
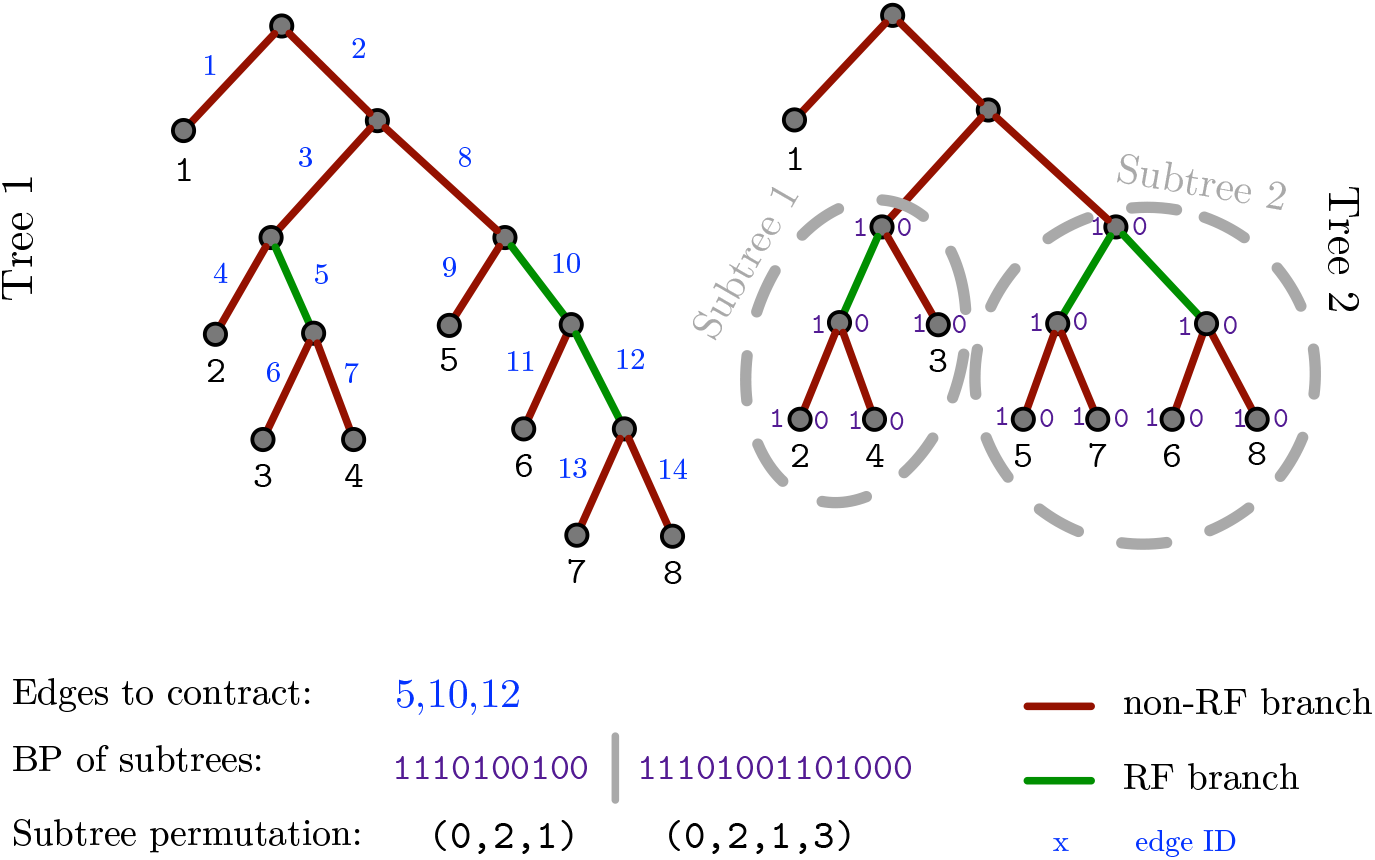
RFC operation example (without branch lengths)

We encode edge contractions by assigning a unique identifier to each branch in DFS order. Then, we store the identifiers of the edges that are contracted in an integer vector. These contraction edges implicitly encode the strict consensus tree.

To construct the second tree, we expand our strict consensus tree again. Since the second tree is also a strictly binary tree, we need to expand each unresolved node in the consensus tree. We do this by first removing all children of an unresolved node. Then, we insert a —previously stored— strictly binary subtree at such an unresolved node. All former children of the unresolved node are contained in this stored binary subtree.

As for SC, each binary subtree that needs to be inserted into the typically non-binary strict consensus tree, is stored in BP representation with an associated taxon identifier permutation. The BP representations of all binary subtrees that we need to insert into the consensus can be stored in a single contiguous bit vector. The respective taxon identifier permutations are also stored in a corresponding single contiguous integer vector.

We provide an example for RFC in Figure 3. For the sake of simplicity, we omitted branch lengths. Edges colored in red are non-RF branches (i.e., part of the consensus tree). Green edges are RF branches that are only present in one of the two trees, but not both. We enumerate the branches in the first tree via a DFS. The vector for storing the edges we contract, contains the identifiers of the green branches (e.g. 5, 10, and 12). The subtrees we insert into the consensus tree to construct the second tree are stored (i) in a BP vector and (ii) a taxon identifier permutation vector. The taxon identifiers of the first subtree have to be mapped from (2, 3, 4) to (2, 4, 3). Therefore, the respective permutation vector is (0, 2, 1). The same applies to the second subtree.

We store branch lengths in two vectors. The first vector stores all RF branch lengths in the order by which they are inserted into the consensus tree. The second vector implicitly stores the branch lengths of the non-RF branches. Thereby, we exploit the property that non-RF branch lengths can be jointly stored for the tree pair when the respective bipartitions are present in both trees. Using this joint storage scheme, we do not store the absolute branch length values of the second tree separately. Instead, we only store the respective branch length difference of the shared bipartitions. Only storing the branch length difference typically requires substantially less space than storing the actual branch length value. This is because branch lengths of successive trees in a MCMC run do generally not change substantially. We can exploit this property by using an appropriate floating point number compression scheme for branch lengths. This scheme requires less space, the smaller the difference between the branch lengths is (see paragraph on branch length compression for details).

To decompress a tree, we are given the preceding tree and our compression data structures. We first contract the edges in the preceding tree via the edge contraction vector to construct the consensus tree. Then, we add the branch length differences to all non-RF branches (i.e., the consensus branches). Subsequently, we traverse the consensus tree via a DFS to identify non-binary nodes. At each non-binary node, we insert the respective binary subtree, as described before. After having inserted a binary subtree at a non-binary node of the consensus tree, our tree is strictly binary and is topologically identical to the second tree. Finally, we set the branch lengths of the newly inserted RF-branches.

#### Implementation

A proof-of-concept SC and RFC implementation in `C/C++` is available at https://github.com/axeltref/tree-compression.git. It uses two external libraries: SDSL and PLL modules. SDSL (Succinct Data Structure Library [9]) offers a plethora of succinct data structures. PLL (Phylogenetic Likelihood Library [10]) modules provides methods for handling phylogenies (e.g., extracting bipartitions, computing RF distances).

Our program takes Newick-formatted trees as input format. The Newick format is supported by widely used phylogenetic inference programs such as RAxML or MrBayes. We provides two tree compression functions: a `simple_compression` and a `rf_distance_compression`. The SC parses a single tree in Newick format, the RFC parses two trees in Newick format. The compressed trees are written to a file that can be decompressed using the `simple_uncompression` and `rf_distance_uncompression` methods, respectively. The decompressed tree is then written to file in Newick format again.

### Branch Length Compression

A core component of our implementation is branch length compression. It transforms double precision floating point numbers, which are typically used internally by phylogenetic inference programs to store branch lengths, to integers and then stores and compresses them appropriately. To achieve this, we initially define the maximum number of decimals *e* we intend to store. Then, we multiply each branch length by 10^e^ and round the result to an integer. Here, we set *e* ≔ 9 to store as many decimals as MrBayes does in its Newick tree files. Next, we search for the maximum among those scaled branch lengths. The number of bits needed to represent this maximum then also suffices to accurately represent all smaller scaled branch lengths. Using SDSL, we construct an integer vector with elements of fixed size (e.g., a vector where each integer element has a fixed size of 11 bits). This fixed size is given by the number of bits needed to represent the maximum scaled branch length. We further compress the fixed-size integer vector via wavelet tree compression. Wavelet trees are succinct data structures for storing strings in compressed form [11].

## Results and Discussion

### Test Datasets

To test our implementation, we calculated MCMC tree samples on empirical DNA datasets containing 500 [12] (denoted as D500) and 354 [13] (denoted as D354) sequences.

### Tree Samples

We generated posterior tree samples for D500 and D354 using MrBayes v. 3.2.6 on a server equipped with an Intel i7-2600 CPU running at 3.40GHz and 16GB RAM. We executed MrBayes for 500, 000 generations and obtained a sample of 1, 000 trees for each dataset.

### Verification

We tested if our algorithm correctly compresses/decompresses trees by applying our SC to every tree and our RFC to every consecutive sample tree pair. We then verified that the decompressed tree was identical to the original tree with respect to (i) our internal tree representation and (ii) with respect to the corresponding Newick tree file.

### Compression Results

The 1000 trees generated by MrBayes are stored in Newick format. The uncompressed size of a single Newick tree file for D500 is 15, 850 Bytes. We applied SC to every tree as well as RFC to consecutive tree pairs (trees (0, 1), (1, 2), etc.). We compared our compression algorithm to state-of-the-art universal compression methods (`bzip2, xy, gzip`). For Newick files, `bzip2` yielded the best compression rate.

Figure 4 shows the resulting compressed tree file sizes for `bzip2` (blue), SC (green), and RFC (red) over the 1, 000 trees for D500.

**Figure 4:**
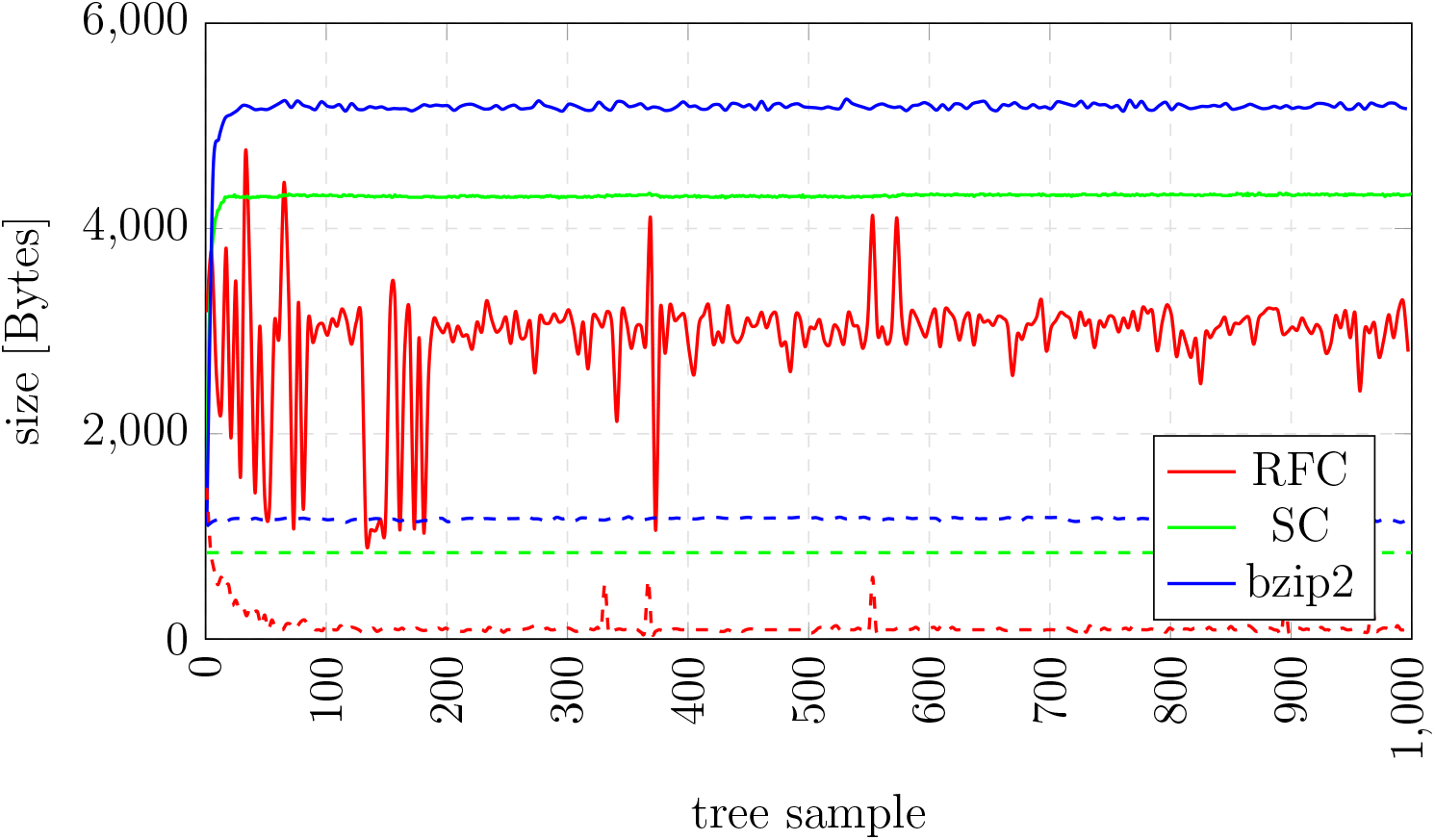
File size of different compression schemes for D500. Solid lines show compression of complete trees, dashed lines show compression of the tree topology.

Averaged over all trees for D500, RFC requires 2, 941 Bytes to store a single tree whereas the best universal compression scheme `bzip2` requires 5, 176 Bytes. Our RFC compression rate is thus 1.76 more efficient than `bzip2` on D500. Note that, the observed variance in the RFC compression rate, especially for the first 500 tree samples, is directly related to the MCMC sampling. As MCMC starts from a random tree, initially a large fraction of topological moves and branch length changes is accepted. As the MCMC procedure moves to areas of higher posterior probability (samples 500–1, 000), the changes to the tree topology and branch lengths become smaller. Thus, the variance in RFC compression rates also decreases.

For D354, RFC requires on average 2, 404 Bytes to store a single tree whereas `bzip2` needs 3, 736 Bytes^1^. Our RFC compression is thus 1.55 times more efficient than `bzip2`.

In Figure 4, we also show the `bzip2`, SC, and RFC compression rates on dataset D500 for tree topologies *without* branch lengths. We applied the methods to a set of modified Newick trees from which we removed the branch lengths via an appropriate script. The average file size for RFC is 121 Bytes while the average size of `bzip2` compressed files is 1, 170 Bytes. Thus, the RFC compression rate is approximately an order of magnitude better than for `bzip2` on this dataset.

Finally, the average compression times for D500 are 6.1 milliseconds for bzip2, 345.2 milliseconds for SC, and 729.1 milliseconds for RFC^1^. Our compression times are about an order of magnitude worse than for `bzip2`. As our implementation has not been optimized yet with respect to compression times, we believe that there is a substantial potential for further improving performance using standard algorithm engineering techniques. In addition, compression times are not performance critical in our context as a phylogenetic tree is only written to file every 1, 000 generations of a MCMC run and the compression can be handled via a separate asynchronous compression thread such that MCMC computations can carry on. Thus our key objective was to optimize the compression ratio and not time.

## Conclusion & Future Work

We presented two novel methods for compressing phylogenetic tree streams as generated by MCMC methods. Our RFC achieves substantially better compression rates than current universal compression schemes such as `bzip2`. When compressing tree topologies without branch lengths, RFC is approximately an order of magnitude better than `bzip2` because the RFC only stores the topological differences between consecutive, relatively similar, MCMC tree samples. Our results also show, that branch lengths clearly constitute the main compression bottleneck. Our RFC might also be applicable to other streams of relatively similar consecutive trees (e.g., XML files have a tree-like structure). One main direction of future work will be to explore, if typical post-analysis steps on the posterior tree distribution, such as building consensus trees or calculating branch length summary statistics, can be directly computed on the compressed data structures.

## Acknowledgements

Part of this work has been funded by the Klaus Tschira Foundation. The authors wish to thank Simon Gog for initial discussions on the algorithm and manuscript.

## Footnotes

1 Additional plots can be found under https://github.com/axeltref/tree-compression/blob/master/plots/phylogenetic_streams_plots.pdf.

